# Identifying candidate *Culicoides* spp. (Diptera: Ceratopogonidae) for the study of interactions with *Candidatus* Cardinium hertigii (Bacteroidetes)

**DOI:** 10.1101/2020.09.21.306753

**Authors:** Jack Pilgrim, Stefanos Siozios, Matthew Baylis, Gert Venter, Claire Garros, Gregory D. D. Hurst

## Abstract

*Culicoides* biting midges (Diptera: Ceratopogonidae) are disease vectors responsible for the transmission of several viruses of economic and animal health importance. The recent deployment of *Wolbachia* with pathogen-blocking capacity to control viral disease transmission by mosquitoes has led to a focus on the potential use of endosymbionts to control arboviruses transmitted by other vector species. Previous screens of *Culicoides* have described the presence of *Candidatus* Cardinium hertigii (Bacteroidetes). However, the biological impact of this symbiont is yet to be uncovered and awaits a suitable system to study *Cardinium*-midge interactions. To identify candidate species to investigate these interactions, accurate knowledge of the distribution of the symbiont within *Culicoides* populations is needed. We used a sensitive nested PCR assay to screen *Cardinium* infection in 337 individuals of 25 *Culicoides* species from both Palearctic and Afrotropical regions. Infections were observed in several vector species including *C. imicola* and the pulicaris complex (*C. pulicaris, C. bysta, C. newsteadi* and *C. punctatus*) with prevalence ranging from low and intermediate, to fixation. Infection in *C. pulicaris* was very rare in comparison to a previous study, and there is evidence the prior record of high prevalence represents a laboratory contamination error. Phylogenetic analysis based on the Gyrase B gene sequence grouped all new isolates within “group C” of the genus, a clade which has to date been exclusively described in *Culicoides*. Through a comparison of our results with previous screens, we evaluate the suitability of *Cardinium*-infected species for future work pertaining to the symbiont.

## Introduction

Worldwide, biting midges of the genus *Culicoides* (Diptera: Ceratopogonidae) are known to transmit more than 50 arboviruses (Mellor, Boorman and Baylis, 2000). Notably, bluetongue virus (BTV) and Schmallenberg virus (SBV) pose a great threat to livestock and wild animal welfare and have caused serious economic damage to the European livestock industry. African horse sickness virus (AHSV) causes a highly lethal disease of equids, with past epizootic outbreaks in Africa, the Middle East and southern Europe. Current control methods of BTV and AHSV rely on vaccines which, due to the rapid emergence and uncertainty of circulating strains, can be ineffective as control mechanisms. Thus, novel initiatives are needed to control the spread of these arboviruses.

The ability of microbial symbionts to alter vectorial capacity in disease vectors is receiving increasing attention (Flores and O’Neill, 2018). For example, the endosymbiont *Wolbachia* leads to a blocking effect of several arboviruses in laboratory-reared mosquitoes (van den Hurk *et al*., 2012). The potential to utilize these effects in wild populations is enabled by *Wolbachia*-induced cytoplasmic incompatibility (CI)-embryo death in mating between infected males and uninfected females. CI is exploited as a mechanism to drive the *Wolbachia* into a population to distribute this virus blocking phenotype into a naïve population. Importantly, this protective phenotype appears to be associated with RNA viruses leading to the potential for symbiont-based biocontrol in major midge-borne pathogens such as BTV, SBV and AHSV.

The symbiont *Candidatus* Cardinium hertigii (Bacteroidetes) has been found to be widely associated with *Culicoides* (Morag *et al*., 2012; Mee *et al*., 2015; Pagès *et al*., 2017). In this study, we focus on *Cardinium* incidence and diversity. Although the biological role of *Cardinium* in biting midges is elusive, data from different host species demonstrate various *Cardinium*-dependent reproductive effects including parthenogenesis-induction (Zchori-Fein *et al*., 2001). CI has also been observed in *Cardinium* in parasitoid wasps (Hunter, Perlman and Kelly, 2003) providing a potential drive mechanism for future introductions into *Culicoides* populations. Importantly, potential effects relating to pathogen-blocking and fitness (e.g. fecundity) remain underexplored.

A key requirement for studying *Cardinium* impacts stems from the need for a model system. Generally, a polymorphically infected population (containing infected and uninfected individuals), is preferable for initiating such investigations as these characteristics offer readily available negative (uninfected) controls. This is particularly important when host species cannot be readily maintained in laboratory cultures and symbiont curing by antibiotic treatment is therefore not feasible. In light of this, we investigated the distribution and prevalence of *Cardinium* infections in *Culicoides* populations from Palearctic and Afrotropical regions, using a nested PCR screen approach. These data are then added and compared to previous screen data to assess what we know about the distribution of *Cardinium* in field populations and the suitability of specific midge species for further investigation pertaining to the symbiont.

## Materials and Methods

337 specimens of 25 *Culicoides* species were collected using light traps from 2007 to 2016 across sites spanning France, South Africa, Sweden and the UK. After collection, all midge specimens were stored in 70% ethanol before sexing and morphological identification. DNA extractions were prepared based on the protocol of Ander *et al*. (2013). In addition, extracts from a previous study by Lewis *et al*. (2014), describing the positive infection status of *C. pulicaris* and *C. punctatus* in the UK, were used to validate screening assays.

Amplification of the host *COI* gene was initially assessed as a means of quality control by conventional PCR assay. DNA extracts which passed quality control were then screened for *Cardinium* using conventional and nested primers amplifying partial sequences of the Gyrase B *(GyrB*) gene (Primer sequences and PCR cycling conditions in Table S1). Collection sites of *Culicoides* populations were then plotted against *Cardinium* infection prevalence based on the nested PCR assay and mapped geographically.

The relatedness of *Cardinium* strains from different host species was analysed using nucleotide sequences from conventional PCR. The Gyrase B gene (*GyrB*) was chosen for phylogenetic analysis because it has a higher level of divergence when compared to the conserved 16S rRNA gene, another gene used in *Cardinium* phylogeny reconstruction. Amplicons were purified enzymatically (ExoSAP) before being sequenced using a BigDye® Terminator v3.1 kit (Thermo Scientific, Waltham, USA), and capillary sequenced through both strands on a 3500 xL Genetic Analyser (Applied Biosystems, Austin, USA). Sequences were aligned using the LINSI algorithm in MAFFT v7 (Katoh and Standley, 2013). A maximum likelihood (ML) phylogeny was then inferred with RAxML v8 (Stamatakis, 2014) using 1000 rapid bootstrap replicates and using the GTR + I + G model which was selected with jModelTest 2 (Darriba *et al*., 2012) using the Akaike information criterion, with the topology search taking the best of Subtree Pruning and Nearest Neighbour Interchange rearranging.

## Results/Discussion

The screening of midge populations in both Palearctic and Afrotropical regions produced varying prevalence of *Cardinium* (Table 1). Most populations did not show any signs of *Cardinium* infection (23/35 populations), five populations showed signs of low-intermediate prevalence, and seven were at fixation for *Cardinium* although a few of these had low sample sizes. This varying prevalence is similar to the study by Mee *et al*. (2015), in which 3/26 positive species had a fixed infection, and six carried *Cardinium* in under 20% of individuals sampled, with the remainder being of variable-intermediate prevalence. However, Mee’s study (2015) also detected at least one *Cardinium* positive individual in each population screened which is at odds to our findings and another study by Pagès *et al*. (2017), investigating *Cardinium* distribution in *Culicoides* from Spain.

**Table 1.** *GyrB* conventional and nested PCR assay results. *Culicoides newsteadi* haplotypes are designated by ^a^Pagès *et al*., 2009, ^b^Ander *et al*., 2013, ^c^*Culicoides newsteadi* N6 previously undesignated. Bold entries are of species identified as being infected with *Cardinium*.

| Subgenus | *Culicoides*<br>Species | Locality<br>(Site name) | Year of<br>collection | Proportion of<br>*Cardinium* positive<br>conventional PCR<br>(n)<br>[95% Confidence<br>interval] | Proportion of<br>*Cardinium*<br>positive nested<br>PCR (n)<br>[95% Confidence<br>interval] |
| --- | --- | --- | --- | --- | --- |
| Avaritia | *C. bolitinos* | South Africa<br>(Onderstepoort) | 2016 | 0 (19) [0-0.21] | 0 (19) [0-0.21] |
|  | *C. huambensis* | South Africa<br>(Koeburg) | 2007 | 0 (1) [0-0.95] | 0 (1) [0-0.95] |
|  | ***C. imicola*** | Corsica (Site 1) | 2015 | 0 (46) [0-0.1] | 0 (46) [0-0.1] |
|  |  | Corsica (Site 2) |  | 0 (27) [0-0.16] | 0 (27) [0-0.16] |
|  |  | **South Africa**<br>**(Onderstepoort)** | **2016** | **0.3 (33) [0.16-0.49]** | **0.3 (33) [0.16-0.49]** |
|  | *C. tuttifrutti* | South Africa<br>(Chintsa) | 2016 | 0 (10) [0-0.34] | 0 (10) [0-0.34] |
|  | *C. obsoletus* | UK (Neston) | 2012-2015 | 0 (33) [0-0.13] | 0 (33) [0-0.13] |
| Beltranmyia | *C. salinarius* | Sweden<br>(Unknown site) | 2009 | 0 (1) [0-0.95] | 0 (1) [0-0.95] |
|  | ***C. sphagnumensis*** | **Sweden**<br>**(Axvalla)** | **2008** | **1 (1) [0.05-1]** | **1 (1) [0.05-1]** |
| Culicoides | ***C. bysta*** | **Corsica (2BPL2)** | **2015** | **0.5 (10) [0.24-0.76]** | **1 (10) [0.66-1]** |
|  | *C. impunctatus* | Sweden (Torsås) | 2008 | 0 (20) [0-0.2] | 0 (20) [0-0.2] |
|  |  | UK (Kielder) | 2016 | 0 (13) [0-0.28] | 0 (13) [0-0.28] |
|  | *C. magnus* | South Africa<br>(Koeburg) | 2007 | 0 (1) [0-0.95] | 0 (1) [0-0.95] |
|  | ***C. newsteadi***<br>**N1<sup>a</sup>** | **Corsica (2APL7)** | **2015** | **0 (4) [0-0.6]** | **0.75 (4) [0.22-0.99]** |
|  |  | **Corsica (2BPL2)** | **2015** | **0 (2) [0-0.8]** | **1 (2) [0.2-1]** |
|  | ***C. newsteadi***<br>**N2<sup>a</sup>** | **Corsica (2BPL2)** | **2015** | **1 (2) [0.2-1]** | **1 (2) [0.2-1]** |
|  | *C. newsteadi*<br>N3<sup>b</sup> | Sweden<br>(Unknown site) | 2008-2010 | 0 (4) [0-0.6] | 0 (4) [0-0.6] |
|  | *C. newsteadi*<br>N6<sup>c</sup> | **Corsica (2APL7)** | **2015** | **0 (12) [0-0.3]** | **0.42 (12) [0.16-**<br>**0.71]** |
|  |  | Corsica (2BPL2) | 2015 | 0 (3) [0-0.69] | 0 (3) [0-0.69] |
|  | *C. pulicaris* | UK (Canterbury) | 2014 | 0 (3) [0-0.69] | 0 (3) [0-0.69] |
|  |  | UK (Hereford) | 2014 | 0 (1) [0-0.95] | 0 (1) [0-0.95] |
|  |  | UK (Luton) | 2014 | 0 (1) [0-0.95] | 0 (1) [0-0.95] |
|  |  | **UK**<br>**(Wolverhampton)** | **2013** | **0 (15) [0-0.25]** | **0.07 (15) [0-0.34]** |
|  |  | UK (Worcester) | 2014 | 0 (7) [0-0.44] | 0 (7) [0-0.44] |
|  | *C. punctatus* | **Sweden**<br>**(Torsås)** | **2008** | **0.44 (25) [0.25-**<br>**0.65]** | **0.8 (25) [0.59-**<br>**0.92]** |
|  |  | **UK (Luton)** | **2014** | **0 (2) [0-0.8]** | **1 (2) [0.2-1]** |
|  |  | **UK**<br>**(Wolverhampton)** | **2014** | **0.4 (5) [0.3-0.99]** | **1 (5) [0.46-1]** |
| Meijerehelea | *C. leucostictus* | South Africa<br>(Kuleni) | 2014 | 0 (1) [0-0.95] | 0 (1) [0-0.95] |
| Monoculicoides | *C. stigma* | Sweden<br>(Unknown site) | 2008 | 0 (3) [0-0.69] | 0 (3) [0-0.69] |
| Oecacta | *C. clastrieri* | Sweden<br>(Unknown site) | 2009 | 0 (2) [0-0.8] | 0 (2) [0-0.8] |
|  | *C. duddingstoni* | Sweden (Bara) | 2008 | 0 (4) [0-0.6] | 0 (4) [0-0.6] |
| Silvaticulicoides | ***C. achrayi*** | **Sweden**<br>**(Axvalla)** | **2009** | **0 (1) [0-0.95]** | **1 (1) [0.05-1]** |
|  | *C. subfascipennis* | Sweden<br>(Romakloster) | 2008 | 0 (12) [0-0.3] | 0 (12) [0-0.3] |
| Silvicola | *C. grisescens* | Sweden (Torsås) | 2008 | 0 (6) [0-0.48] | 0 (6) [0-0.48] |
| Synhelea | *C. similis* | South Africa<br>(Alexandria) | 2016 | 0 (6) [0-0.48] | 0 (6) [0-0.48] |
| Wirthomyia | *C. reconditus* | Sweden<br>(unknown site) | 2008 | 0 (1) [0-0.95] | 0 (1) [0-0.95] |

It is possible that the apparent “hotspot” in Australia is as a result of a more sensitive assay, as the qPCR method used by Mee et al. (2015) is sometimes preferred to nested PCR in screening. Alternatively, variation in thermal environment could explain the prevalence discrepancies observed between these bioclimatic zones. Morag *et al*. (2012) found a positive correlation between land surface temperature (LST) and *Cardinium* prevalence in *Culicoides imicola* in Israel, with higher prevalence at higher mean LST. Furthermore, there is a known relationship between *Cardinium* presence and latitude more globally, with *Cardinium* infection being more common for hosts near the equator (Charlesworth *et al*., 2019).

In order to investigate *Cardinium*-*Culicoides* interactions, candidate model midge species must be identified, which combine vectorial capacity with either lab culturability or a natural polymorphism in symbiont presence that permits comparison between infected and uninfected individuals. The conventional assay detected *Cardinium* in four putative vector species of bluetongue virus (BTV); *C. imicola, C. newsteadi, C. bysta* and *C. punctatus* (Table 1). Additionally, evidence of a low-level *Cardinium* infection was detected in one individual of the vector species *C. pulicaris* when screened with the nested assay. The infection patterns within the pulicaris complex species (*C. bysta, C. newsteadi, C. pulicaris* and *C. punctatus*) are of particular intrigue due to conflicting reports of *Cardinium* prevalence within the group (Lewis *et al*., 2014; Pagès *et al*., 2017). Notably, our more sensitive nested PCR assay detected a very low *Cardinium* prevalence of 3.7% (1/27) for *C. pulicaris*, compared to Lewis *et al*. where a moderate prevalence of 34.5% (10/29) was described. However, when rescreening Lewis et al.’s DNA extracts, the COI barcode vouchers from “*C. pulicaris*” *Cardinium* positives matched *C. punctatus* indicating cross-contamination of DNA extractions had occurred and the infection detected previously must be considered artefactual (Details in supplemental file 1). This observation also explains why the *GyrB* sequences deposited for both *C. pulicaris* and *C. punctatus Cardinium* strains are identical (Accession number: HG380244). Overall, our findings, together with the observations of Pagès et al. (describing 0/606 *C. pulicaris* infections), suggests *C. pulicaris Cardinium* infection is not common and likely has little biological significance. Despite this, a species from the pulicaris complex (*C. bysta;* Corsica) contained *Cardinium* in all individuals tested (Table 1) suggesting that distinguishing cryptic species through molecular markers is important when assessing candidates for investigating symbiont interactions.

Phylogenetic analysis based on a 1300 bp region of the *GyrB* gene confirmed a monophyletic clade of *Cardinium*, grouping all *Culicoides Cardinium* isolates from this study in group C of the genus, in which all previously described sequences of *Culicoides* clustered (Figure 1) (Accession numbers: LR877462-LR877465). The *C. imicola GyrB* sequence obtained from South Africa in this study (Accession number: LR877465) was identical to both of those obtained from Kenya (Accession number: KR026927) and Israel (Accession number: JN166963). Likewise, the *Cardinium* sequences obtained from *C. punctatus* from Wolverhampton, UK (Accession number: LR877464) had 100% identity to the sequence reported in the same species by Lewis *et al*. (Accession number: HG380244). Mee *et al*. (2015) has previously suggested that *Cardinium* strains group by geography; in contrast, we find a sporadic distribution of strains with respect to location. For example, the *C. bysta* and *C. newsteadi* strains (Accession numbers: LR877462 and LR877463) identified from Corsica group with strains from Israel and Japan respectively (Accession numbers: JN166964 and AB506792). Furthermore, no clear pattern of *Cardinium* distribution was observed between the three geographical regions of this study (Fig. S1).

**Figure 1.**
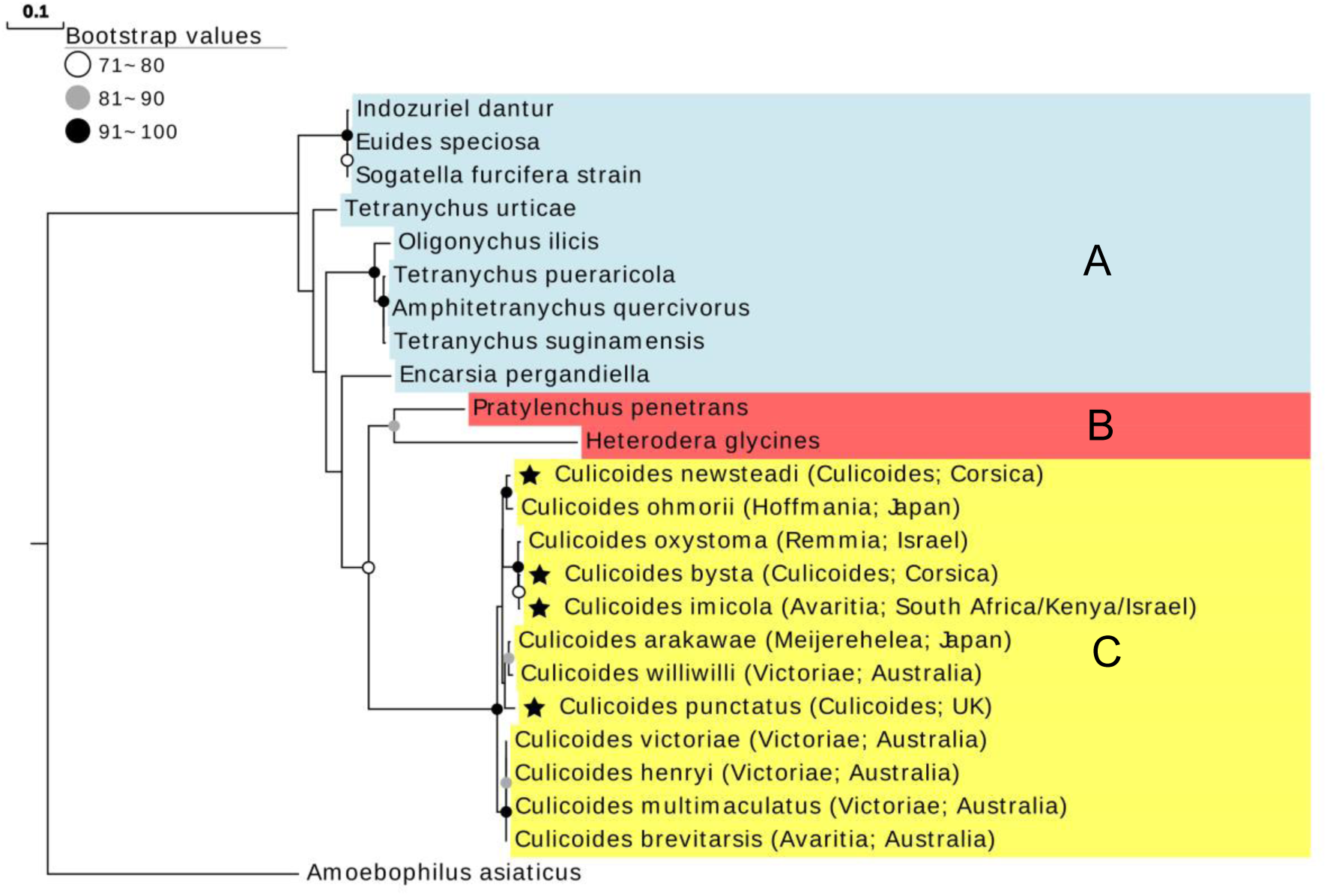
A maximum likelihood phylogeny based on a 1300 bp region of the *GyrB* gene generated in RAxML using 1000 rapid bootstrap replicates and using the GTR + I + G substitution model. Groups A (blue) B (red) and C (yellow) are designated based on *Cardinium* taxonomic convention. Stars represent sequences generated in this study. Brackets indicate subgenus and country of origin.

In contrast with other symbiont-insect associations, the biological significance of *Cardinium* in *Culicoides* remains unknown and requires further research. Despite a likely vertical transmission route, our observed lack of sex-bias in infection observed in this study (*C. punctatus*; Fisher’s exact, *p*=0.76 and *C. imicola*; Fisher’s exact, *p*=0.68) corroborates previous studies (Morag *et al*., 2012; Mee *et al*., 2015; Pagès *et al*., 2017) and suggests the induction of parthenogenesis, feminisation or male-killing is unlikely to be associated with *Cardinium*. Other *Cardinium* strains have been implicated in cytoplasmic incompatibility (CI), including in the wasp *Encarsia pergandiella* (Hunter, Perlman and Kelly, 2003). Our recent publication of the *C. punctatus Cardinium* genome by Siozios *et al*. (2019) has suggested possible unique genes related to CI. Overall, these observations suggest potential *Cardinium*-induced CI should be a priority of investigation the future due its possible role in driving the symbiont (and any desired effects) into midge populations.

The major Palearctic vector of BTV, *Culicoides obsoletus*, has previously been shown to contain *Cardinium* at 0.4% prevalence (Pagès *et al*., 2017). We did not find the symbiont in our screen. A low prevalence indicates *Cardinium* is likely not relevant to vector biology at a population level and is not a suitable candidate for investigation. However, two other important vector species, *Culicoides imicola* and *Culicoides sonorensis*, have been reported to carry *Cardinium*-infection (Table 1; Morag *et al*., 2012; Möhlmann, 2019). Thus, these species appear to be the most promising candidate species for investigating *Cardinium* effects on vectorial capacity. As certain *C. imicola* and *C. sonorensis* populations are polymorphically infected, this allows for the assessment of naturally occurring negative controls without the need for the confounding effects of curing through antimicrobials. In favour of *C. imicola* as a model is the ease of obtaining large field catches, but barriers in laboratory cultivation are still to be overcome. However, *C. sonorensis* laboratory colonies already exist, suggesting this species is the most promising candidate for investigating symbiont-virus interactions and symbiont-mediated reproductive effects.

## Notes

### Competing Interest Statement

The authors have declared no competing interest.

