## Supplemental File 1 for "Identifying candidate *Culicoides* spp. (Diptera: Ceratopogonidae) for the study of interactions with *Candidatus* Cardinium hertigii (Bacteroidetes)"

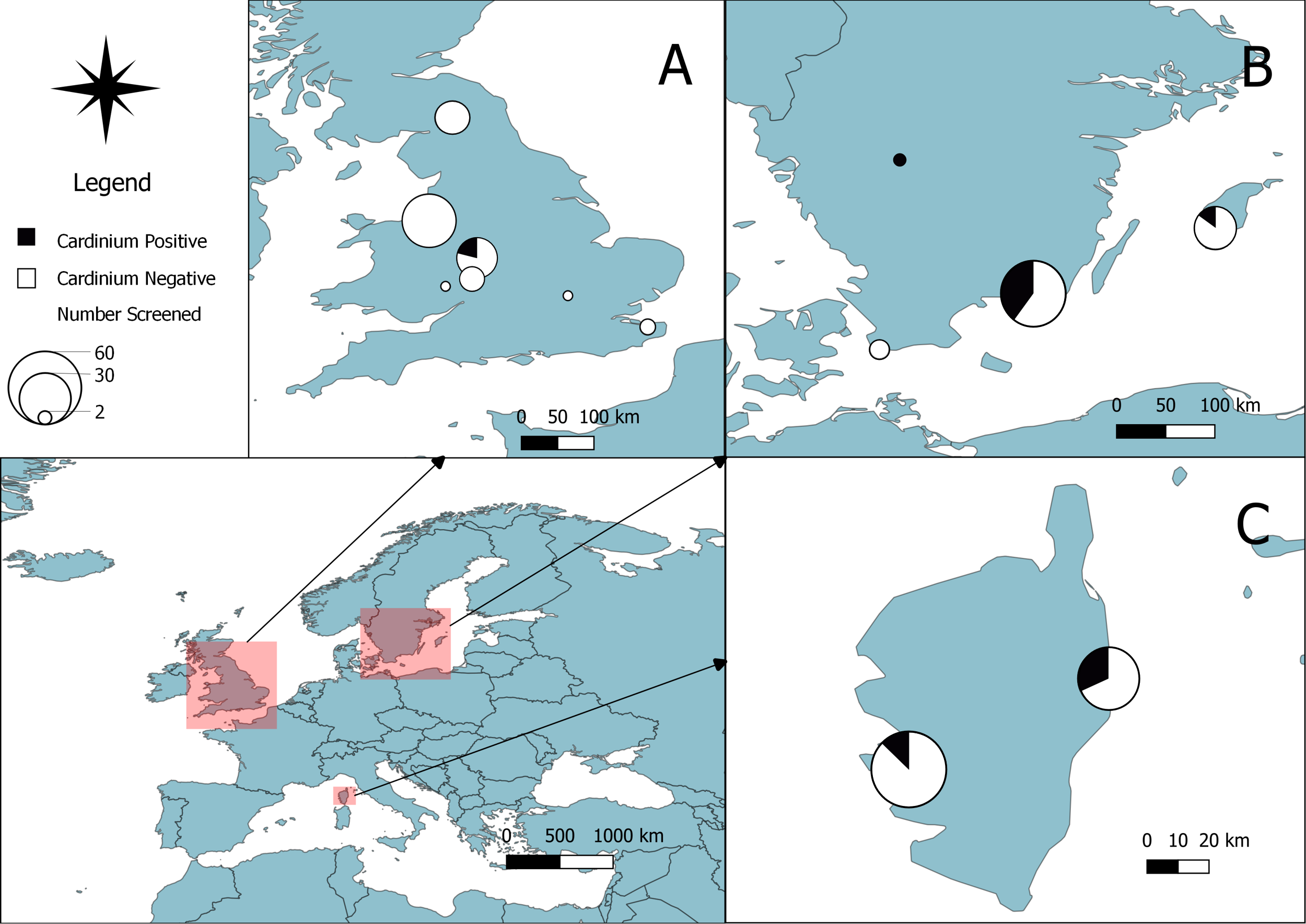

**Figure S1.1.** Palearctic *Culicoides* trapping and endosymbiont distribution. A QGIS map depicting collection sites across the UK (A), Sweden (B), and Corsica, France (C). The size of pie charts represents the number of individuals screened, with the proportion of *Cardinium*-infected *Culicoides* designated in black. White circles indicate individuals where no *Cardinium* was detected by *GyrB* conventional or nested PCR.

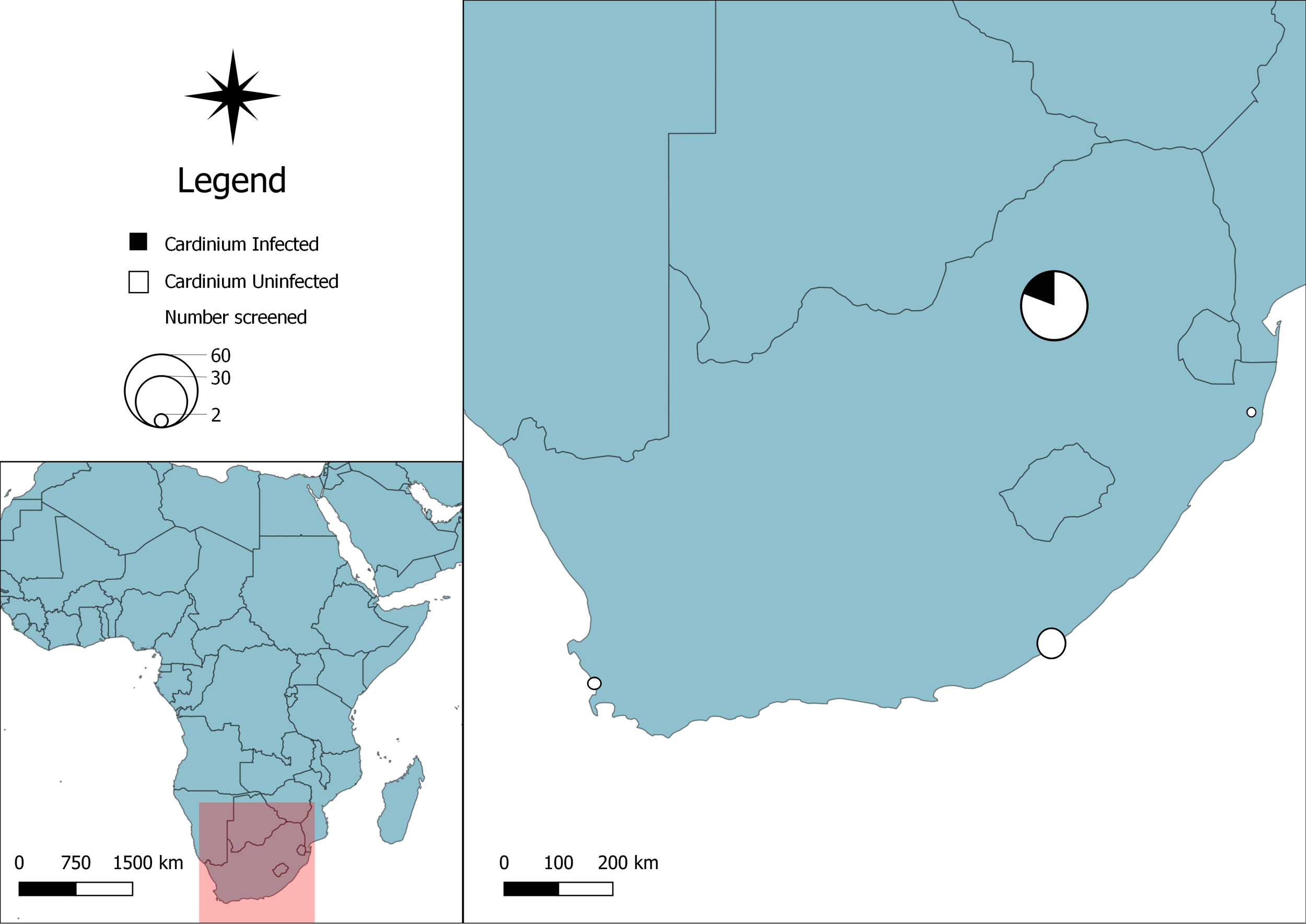

**Figure S1.2** Southern Afrotropical *Culicoides* trapping and endosymbiont distribution. A QGIS map depicting collection sites across four sites in South Africa. The size of pie charts represents the number of individuals screened, with the proportion of *Cardinium*-infected designated in black. White circles indicate individuals where no *Cardinium* was detected by *GyrB* conventional or nested PCR.

**Table S1.1.** *COI* and *Gyrase B* gene primer attributes.

| **Target** | **Primer name** | **Sequence (5’-3’)** | **Tm (°C)** | **Size (bp)** | **Reference** |
| --- | --- | --- | --- | --- | --- |
| Cytochrome oxidase subunit 1 (*COI*) | LCO2198 | GGTCAACAAATCATAAAGATATTGG | 51 | 708 | Folmer *et al.* 1994 |
|  | HCO1490 | TAAACTTCAGGGTGACCAAAAAATCA | 56 |  |  |
|  | C1‐J‐1718 | GGAGGATTTGGAAATTGATTAGT | 52 | 523 | Dallas *et al.* 2003 |
|  | C1‐N‐2191 | CAGGTAAAATTAAAATATAAACTTCTGG | 50 |  |  |
| Gyrase B | gyrB23F | GGAGGATTACATGGYGTGGG | 60 | 1368 | Lewis *et al.* 2014 |
|  | gyrB1435R | GTAACGCTGTACATACACGGCATC | 60 |  |  |
|  | gyrBnest212F | AAGGCAACCCTATGCACCAA | 59 | 347 | This study |
|  | gyrBnest654R | GGYCTTAGTTTGCCCTTCAAATTG | 59 |  |  |

**Table S1.2.** PCR cycling conditions

| **Target** | **Initialisation** | **Denaturation, annealing, extension** | **Final extension, hold** | **Number of cycles** |
| --- | --- | --- | --- | --- |
| Cytochrome oxidase subunit 1 (COI) | 95°C/5min | 95°C/30s, 50°C/1min, 72°C/1min | 72°C/7min, 15°C/∞ | 35 |
| Gyrase B (Conventional) | 95°C/5min | 95°C/30s, 55°C/1min, 72°C/1min30s | 72°C/7min, 15°C/∞ | 35 |
| Gyrase B (Nested) | 95°C/5min | 95°C/25s, 55°C/30s, 72°C/45s | 72°C/7min, 15°C/∞ | 35 |

*Clarification of the Cardinium infection status of C. pulicaris and C. punctatus from Lewis et al. study*

Of the 26 *C. punctatus* DNA extracts from Lewis’ study screened, 25/26 (96%) were positive for *Cardinium*, all of which were validated for sufficient DNA quality via PCR amplification of the *COI* subunit using the HCO1490/LCO2198 primer set. Despite adequately replicating Lewis’ findings for *C. punctatus*, only 12/39 (31%) *C. pulicaris* samples were positive when using the HCO1490/LCO2198 primer pair, as opposed to 39/39 (100%) claimed in the previous study. Incidentally, the 12 *COI* positive samples were the same DNA extracts which came back PCR positive for the *Cardinium* *GyrB* screen as documented by Lewis. Furthermore, fresh *C. pulicaris* caught in the autumn of 2015 did not show positive signals for amplification of *COI* using HCO1490/LCO2198 primers. This led to the likelihood that there was a polymorphism at one of the forward or reverse primer sites designed to amplify *COI* in *C. pulicaris*. To test this, a new *COI* locus was chosen to observe whether the previous negative samples would become positive. The secondary quality control primers C1‐J‐1718/C1‐N‐2191 amplified all the *C. pulicaris* DNA extracts confirming this premise.

Although all of these samples had now been confirmed to have adequate DNA quality, there were still the unresolved issues of how Lewis had detected *COI* positive *C. pulicaris* with the invalid HCO1490/LCO2198 primers, and why the re-screened HCO1490/LCO2198 positives were coincidentally the only samples to come back positive for *Cardinium*. Obtaining the raw data from Lewis’ study resolved the first of these issues. Examination of Lewis’ gel images corroborated my re-screening data (see below); 12/39 *C. pulicaris* samples gave positive bands after amplification with HCO1490/LCO2198 primers despite Lewis claiming that all passed the *COI* quality control screen. To explain the matter of why there were any HCO1490/LCO2198 amplified *C. pulicaris* extracts to begin with, there were several possible explanations: the DNA extracts had been cross contaminated with other *Culicoides*’ extracts through a pipetting error; the individuals had been misidentified when morphologically distinguishing the *Culicoides* species and were in fact another species*;* or the individuals morphologically appeared as *C. pulicaris* but were an unidentified cryptic species which lack the polymorphism at the HCO1490/LCO2198 site and are infected with *Cardinium*. To distinguish between these possibilities, Sanger sequencing of the purified PCR product of both HCO1490 and C1-N-2191 was undertaken and compared to known mitochondrial DNA barcodes on Genbank. BLASTn alignment identified the HCO1490 product to be 99% homologous to *C. punctatus* COI (Accession number: KX064676), whereas the C1-N-2191 product was 100% identical to *C. pulicaris* COI (Accession number: KJ624116). As both loci overlap with each other, these data implied cross contamination of *C. pulicaris* and *C. punctatus* DNA extracts (See below).

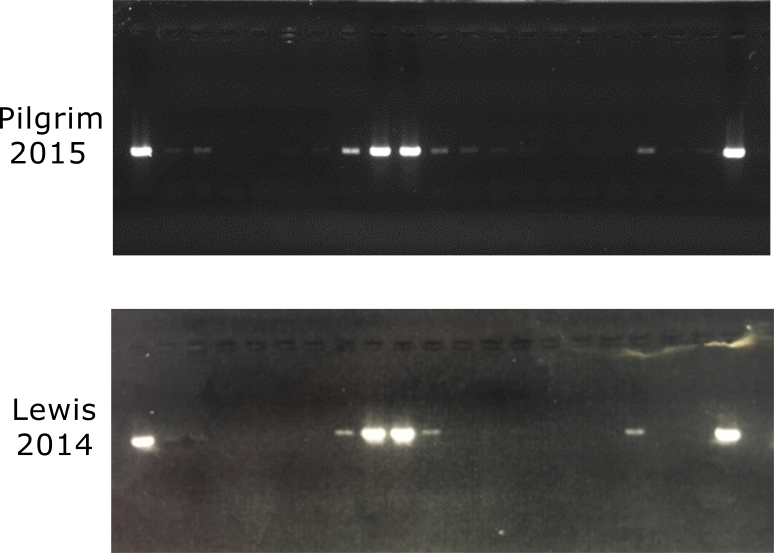

Quality control of DNA extracts labelled as *Culicoides pulicaris* from Lewis *et al*.’s study using HCO1490/LCO2198 primers (Folmer *et al.* 1994). Gel images are taken from a screen undertaken separately by the author in this study (2015) and by Lewis *et al.* (2014).

| **Identifier** | **LCO1490/HCO2198 (Folmer *et al.* 1994)** | **C1J-1718/C1N-2191 (Dallas *et al.* 2003)** | ***Cardinium* infection** |
| --- | --- | --- | --- |
| Lewis | 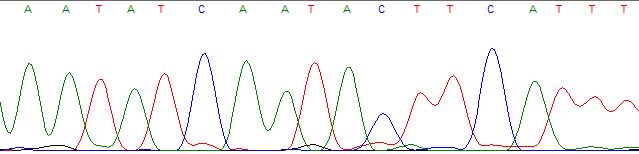 | 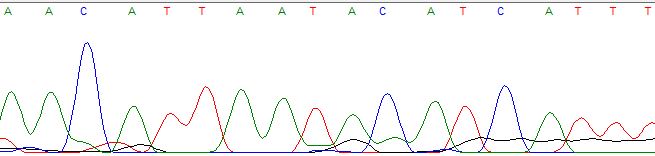 | + |
|  | >99% homology Culicoides punctatus (KX064676) | >99% homology Culicoides pulicaris (KJ624116) |  |
| Pilgrim | N/A | 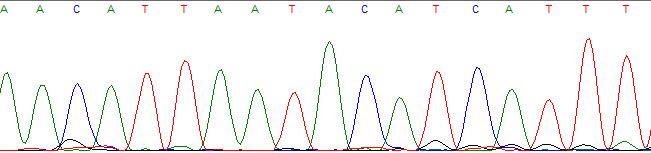 | - |
|  |  | >99% homology Culicoides pulicaris (KJ624116) |  |

Sequencing of *COI* amplicons from samples initially identified as *Culicoides pulicaris* from Lewis *et al.* (2014) and this study, suggesting cross-contamination with *Culicoides punctatus* in *Cardinium* positives from Lewis’ study. N/A=Non-amplifiable.

| ***Culicoides* species** | **LCO1490/HCO2198 (Folmer *et al.* 1994)** | **C1J-1718/C1N-2191 (Dallas *et al.* 2003)** |
| --- | --- | --- |
| *C. pulicaris* | - | + |
| *C. bysta* | + | + |
| *C. punctatus* | + | - |
| *C. obsoletus* | + | + |
| *C. impunctatus* | + | + |

PCR amplification status of *Culicoides* species of interest with two sets of *COI* primers.
